# Structural basis of DNA packaging by a ring-type ATPase from an archetypal viral system

**DOI:** 10.1101/2022.05.10.491410

**Authors:** Herman K.H. Fung, Shelley Grimes, Alexis Huet, Robert L. Duda, Maria Chechik, Joseph Gault, Carol V. Robinson, Roger W. Hendrix, Paul J. Jardine, James F. Conway, Christoph G. Baumann, Alfred A. Antson

## Abstract

Many essential cellular processes rely on substrate rotation or translocation by a multi-subunit, ring-type NTPase. A large number of double-stranded DNA viruses, including tailed bacteriophages and herpes viruses, use a homomeric ring ATPase to processively translocate viral genomic DNA into procapsids during assembly. Our current understanding of viral DNA packaging comes from three archetypal bacteriophage systems: *cos, pac* and phi29. Detailed mechanistic understanding exists for *pac* and phi29, but not for *cos*. Here we reconstituted *in vitro* a *cos* packaging system based on bacteriophage HK97 and provided a detailed biochemical and structural description. We used a photobleaching-based, single-molecule assay to determine the stoichiometry of the DNA-translocating ATPase large terminase. Crystal structures of the large terminase and DNA-recruiting small terminase, a first for both this phage and a *cos* system, reveal unexpected mechanistic similarities between *cos* and *pac* systems. At the same time, mutational and biochemical analyses indicate a new regulatory mechanism for ATPase multimerization and coordination in the HK97 system. This work therefore establishes a framework for studying the evolutionary relationships between ATP-dependent DNA translocation machineries in double-stranded DNA viruses.

## INTRODUCTION

Homomeric ring nucleoside triphosphatases (NTPases) are a diverse class of enzymes that activate upon oligomerization to drive important cellular processes such as DNA replication and protein degradation through mechanical movements coupled to nucleoside triphosphate (NTP) hydrolysis. Viral packaging ATPases represent an unusual family of homomeric ring NTPases. They feature a unique set of arginine fingers for inter-subunit coordination and, depending on the virus, they act either as pentamers or hexamers (1–4). Essential for tailed bacteriophage and herpes virus replication, viral packaging ATPases translocate viral genomic DNA into newly assembled, empty procapsids during replication inside the host (5, 6). The packaged DNA is highly compacted and stressed within the viral capsid (7). Hence, a well-coordinated ATPase motor is necessary to ensure processivity and speed to produce a maximum number of viable virus particles during infection. The structure and mechanics of viral packaging ATPases have been elucidated for two of three archetypal packaging systems, *pac* and phi29-like. The third system, *cos*, though biochemically defined based on extensive work on lambda (8), remains structurally uncharacterised, which has prevented complete understanding of this classical viral model system. Consequently, it is still unclear whether a universal assembly mechanism exists among packaging ATPases and whether their regulatory mechanisms are conserved throughout double-stranded DNA viruses that employ them for virus assembly.

*Cos* packaging systems rely on precise cleavage of newly replicated viral DNA, produced as a concatemer of multiple genome copies, to package successive unit-length genomes into procapsids (8). By contrast, *pac* systems feature imprecise, sequence-independent cleavage of the DNA concatemer during head filling, giving rise to variable genome lengths inside virus particles. Both systems encode a large terminase (TerL) protein, composed of a viral packaging ATPase domain and a nuclease domain, and a small terminase (TerS) protein, which recognises the *cos* or *pac* site between genome copies and, upon binding of DNA, recruits TerL to make an initiating DNA cut (9, 10). To enable translocation into procapsids, the newly cut DNA is brought to the portal complex at a unique vertex of the procapsid, where TerL oligomerizes to form an active motor. Once approximately one genome length has been packaged, the TerL nuclease domain is proposed to reactivate, freeing the DNA for packaging into the next procapsid. For *cos* systems, this cleavage event occurs specifically at the next *cos* site (11, 12), whereas for *pac* systems, the cleavage is not specific (13, 14).

The structural details of TerL and TerS are known only for *pac* systems. Crystal structures suggest that *pac* TerS proteins form circular assemblies of 9–11 subunits with helix-turn-helix motifs arrayed on the outside (15–17). *Pac* TerL proteins have been shown to oligomerize on the portal of procapsids, and important residues for mechanochemical coupling have been biochemically identified. However, the stoichiometry of interaction between TerS and TerL is unknown. Analytical ultracentrifugation-based measurements of *cos* phage lambda proteins suggest that TerS subunits pair with one TerL during initial assembly of the motor machinery. Despite *cos* proteins having similar domain predictions, owing to the complete lack of structural information on them, it remains unclear whether motor assembly mechanisms are conserved between the *cos* and *pac* packaging systems. Compositionally distinct from *cos* and *pac* systems, phi29 systems package pre-formed, unit-length, protein-capped genomes (18). Instead of TerL and TerS, they encode a single-domain viral packaging ATPase, which complexes with a structured RNA element (pRNA) on the portal of procapsids to selectively package viral DNA (19). Optical tweezers experiments showed that the ATPase pentamer translocates DNA in bursts of four 2.5-bp steps followed by a long dwell phase at speeds up to 165 bp/s (7, 20). Cryo-EM structures of the DNA-translocating assembly combined with further single-molecule studies have identified two *trans*-acting residues that participate in nucleotide exchange and catalysis, the latter acting like an arginine finger, which stabilised the leaving phosphate of ATP in an adjacent subunit during DNA translocation (1, 21, 22). Indeed, the establishment of a defined *in vitro* packaging system was instrumental in the deriving of mechanistic understanding from this system (19). Similar *in vitro* packaging systems have been built for *cos* phage lambda, and *pac* phages T4 and P23-45, which enabled progressive mapping of the viral DNA packaging reaction and identification of distinct or similar ATPase regulatory elements (2, 23–26).

The *cos* bacteriophage HK97 shares sequence homology with lambda in their tail fibre and proteins involved in transcriptional regulation, DNA replication, integration and lysis (27). Sequence homology in the structural proteins such as the capsid protein, TerL and TerS is low, but the proteins are identifiable based on gene order and ATPase Walker motifs. As one of the classical viral model systems, the capsid structure of HK97 and its assembly and maturation mechanisms have been extensively characterised (28). The *cos* site delineating genome boundaries is also defined (27). Based on this prior work, we established *in vitro* the DNA packaging system of bacteriophage HK97 with the aim of elucidating the general mechanisms of packaging in *cos* viruses. Our *in vitro* system consists of the portal-containing procapsid, the HK97 TerL protein GP2, TerS protein GP1, and DNA substrate containing the *cos* site. We showed biochemically that TerS facilitates packaging arrest at the *cos* site, contributing to specificity in the termination of packaging. By X-ray crystallography and complementary stoichiometry analyses, we discovered that the HK97 TerS and TerL have more in common structurally with their *pac* homologues than previously thought. Finally, we identified by extensive mutagenesis an unusual lysine residue that is critical for ATPase activity, possibly defining a new subclass of viral packaging ATPases among homomeric ring NTPases.

## MATERIAL AND METHODS

### Cloning of HK97 TerL, TerS and *cos* DNA

The coding sequences of TerL and TerS (residues 1–526 and 2–161, respectively) were amplified from CsCl-purified wild-type HK97 phage particles and cloned into pET22a-based vectors using the ligation-independent In-Fusion Cloning system (Clontech) to generate N-terminal His-SUMO fusions and constructs with a short non-cleavable N-terminal His-tag (MGSSHHHHHH). A construct containing a short N-terminal His-tag, followed by emerald GFP with mutations S65T, F64L, S72A, N149K, M153T and I167T, a 10-residue G/S linker, and finally large terminase was generated by insertion of the GFP and linker coding sequence into the His-fusion construct via In-Fusion. Site-directed mutagenesis of the His-tagged TerL construct was performed using an adapted Quick Change protocol (29). The HK97 *cos* region was recovered by extracting a 784-bp segment (−312 to +472) around the *cos* cleavage site by overlap-extension PCR using wild-type phage particles, and cloned using *Bam*HI and *Eco*RI sites into a pUC18 plasmid. All primers used are listed in Table S5.

### Overproduction and isolation of recombinant TerL, TerS and GP74

Recombinant proteins were overproduced using *Escherichia coli* BL21(DE3) pLysS cells in Lysogeny Broth (LB, Miller formulation) containing 30 μg/ml kanamycin and 33 μg/ml chloramphenicol. Expression was induced at OD_600_ = 0.8 with 0.4 mM IPTG at 37 °C for 4 h for TerS and at 16 °C for 18 h for TerL. For selenomethionine labelling, cells were grown in LB with amino acids added at OD_600_ = 0.6 to suppress methionine biosynthesis as described by Van Duyne et al. (30) and L-selenomethionine added to 200 μg/mL concentration on induction with IPTG.

Cells were harvested by centrifugation and lyzed by sonication in 20 mM Tris-Cl, 1 M NaCl, 10% (v/v) glycerol, 20 mM imidazole, 0.05% (v/v) β-mercaptoethanol, pH 8.0 with 100 μM 4-(2-aminoethyl)benzenesulfonyl fluoride (AEBSF), 1 μM leupeptin, 1 μM pepstatin A and 10 μg/mL RNaseA. The soluble fraction after centrifugation was applied to a HisTrap FF Ni Sepharose column (GE Healthcare) and the protein eluted with a 20–500 mM imidazole gradient. Eluate was dialyzed against 20 mM Tris-Cl, 200 mM NaCl, 1 mM dithiothreitol (DTT), pH 8.0, at 4 °C overnight, with 1:100 (w/w) SUMO protease if necessary for untagged protein production. Further purification was performed using a 200–1000 mM NaCl gradient on a cation exchange Mono S column and anion exchange Mono Q column (Amersham) for TerS and TerL, respectively, and finally using Superdex 200 and Superdex 75 16/600 size exclusion columns (GE Healthcare). Proteins were concentrated using Vivaspin centrifugal concentrators (Sartorius) with MWCOs 3000 and 30,000, respectively. Proheads were produced by infection of *Escherichia coli* 594 cells with HK97 amber mutant amC2, propagated using *Escherichia* LE392 cells, and purified by PEG precipitation, glycerol gradient ultracentrifugation and cation exchange chromatography, as previously described (31). GP74 was overproduced using *Escherichia coli* BL21(DE3) pLysS cells, and was purified by Ni-NTA, anion exchange and size exclusion chromatography as described previously (32).

### DNase protection-based packaging assay

HK97 genomic DNA was isolated from CsCl-purified wild-type phage particles by phenol-chloroform extraction and ethanol precipitation. Linear plasmid DNA was prepared using FastDigest restriction endonucleases (Thermo Scientific). DNA packaging was assayed for using an established DNase protection assay (33). In 20 μL packaging buffer (20 mM Tris-Cl, 10 mM MgSO_4_, 30 mM potassium glutamate, 1 mM β-mercaptoethanol), 0.5 μg DNA (2 nM, 13.5 nM and 15 nM for HK97 genomic DNA, linear pUC18 and *cos*-containing pUC18 DNA, respectively) was mixed with procapsids in a 1:2 molar ratio, TerL and TerS at 1 μM and 2 μM monomeric concentrations, respectively, incubated for 5 min at room temperature (20–22°C), and ATP (Sigma-Aldrich) added to 1 mM concentration to initiate packaging. After 30 min at room temperature, unpackaged DNA was digested by incubation with 1 μg/mL DNase I (Roche) for 10 min. DNase was inactivated and packaged DNA was liberated from capsids by incubation with 25 mM EDTA, pH 8.0, 500 μg/mL proteinase K (Roche) at 65°C. Finally, the packaged DNA was analyzed by agarose gel electrophoresis.

### Measurement of ATPase reaction kinetics

ATPase activity was measured using the EnzChek Phosphate Assay Kit (Molecular Probes). Reactions contained 1 μM TerL in packaging buffer, with or without 25 nM *Sca*I-linearised *cos*-containing pUC18, 25 nM procapsid and 2 μM TerS.

### Structure determination by X-ray crystallography

Crystals of His-tagged TerL protein were obtained with streak-seeding by hanging drop vapor diffusion using 20 mg/mL protein solution in 250 mM NaCl, 5% (v/v) glycerol, 2 mM DTT, 20 mM MgSO_4_, 10 mM AMP-PNP, 20 mM HEPES pH 7.5, equilibrated against reservoir containing 1.6 M ammonium sulfate, pH 7.6. To better resolve density for the nuclease domain, the native crystals were soaked in 1.8 M ammonium sulfate, 0.5 M KCl, 10 mM MgSO_4_, 5 mM AMP-PNP, 20% (v/v) glycerol before vitrification. Crystals of selenomethionine-labelled TerL were obtained against 1.5 M ammonium sulphate pH 7.6, by streak-seeding first with native crystals, then with its own crystals, and finally cryo-protected in 5 mM MgSO_4_, 4 mM AMP-PNP, 1.9 M ammonium sulfate, 21% (v/v) glycerol, 10 mM HEPES pH 7.5. Crystals of TerS (22 mg/mL in 300 mM potassium glutamate, 20 mM HEPES pH 7.0) were obtained by hanging drops with reservoir containing 9% (w/v) PEG 3350, 0.1 M succinic acid pH 7.0. For experimental phasing, crystals were soaked in solution containing 10 mM HEPES pH 7.0, 50 mM succinic acid pH 7.0, 18% (w/v) PEG 3350, 15 % (v/v) glycerol, 500 mM KI. A quick pass in the same cryo-protectant but with 250 mM KI did not produce useful anomalous signal but improved the quality of diffraction to 1.4 Å. Data were collected at 100 K at Diamond Light Source beamlines I02 and I03, integrated and scaled with XDS (34). Initial phasing was performed with SHELX (35) by MAD and SAD for TerL and TerS, respectively. Density modification and auto-tracing were performed using SHELXE, followed by iterative cycles of model building and refinement in BUCCANEER and REFMAC (36, 37). Models were completed in COOT (38) and refined against the higher-resolution native datasets. Despite the presence of AMP-PNP and magnesium during the crystallization of TerL, no electron density corresponding to either was observed. Molecular graphics were generated using PyMol (Schrödinger) and UCSF Chimera (39). Electrostatic potentials were calculated using APBS (40) under the SWANSON force field (41).

### Single-molecule total internal reflection fluorescence microscopy-based subunit counting

DNA for motor assembly and surface attachment was prepared by amplifying a 230-bp DNA segment across the HK97 *cos* cleavage site (−80 to +150) using unmodified forward and 5′-biotinylated reverse primers. Imaging sample chambers were constructed by applying 25 μL 1 mg/mL biotin-BSA (Sigma) in PBS with 0.1 mg/mL 5-μm diameter silica beads (Bangs Laboratories) to a quartz slide (UQG Optics) and covering with a washed coverslip (No. 1, 22 mm × 64 mm, Menzel-Gläser), then sealing with nail varnish over the short sides to create a flow cell. After 10 min, unbound biotin-BSA was washed out and the flow cell was equilibrated with two volumes of imaging buffer: 20 mM Tris-Cl, 10 mM MgSO_4_, 30 mM potassium glutamate, 1 mM ATP-γ-S, 0.1 mM β-mercaptoethanol, 0.5 mM Trolox (Sigma-Aldrich), 0.1 mg/mL acetylated BSA (Sigma-Aldrich), 0.25% (w/v) PEG 6000 (Santa Cruz Biotech), 380 nM BOBO-3 stain (Thermo Fisher). 100 nM DNA was incubated with 3.7 μM BOBO-3 stain (Thermo Fisher), 100 nM streptavidin tetramer, 200 nM proheads, 1 μM GFP-TerL and 2 μM TerS for 20 min at room temperature. Adding ATP to 1 mM, the mixture was immediately chased with 10 volumes of imaging buffer. 25 μL of this diluted mixture was drawn into the flow cell and the long sides sealed subsequently with nail varnish. The sample was visualized by prism-coupled TIRFM on a modified inverted IM35 microscope (Carl Zeiss AG). Fluorophores were excited with 488-nm and 561-nm lasers (Coherent) operating at 10 mW and 30 mW, respectively. Incident 488-nm light was circularly polarised using a 488-nm quarter-wave plate (Edmund Optics) to minimise orientation-dependent excitation. Fluorescence emission was captured through a Plan-Apochromat 100×/NA 1.4 oil-immersion objective (Carl Zeiss AG). A dual-view image splitter (OptoSplit II, Cairn Research) with 1.6× magnification, 580 nm long-pass emission dichroic (Zeiss) and bandpass filters for GFP (ET525/50M, Chroma) and BOBO-3 (ET605/20M, Chroma) was used to view the image in two fluorescence emission channels. Video data were recorded using an Evolve 512 electron-multiplying CCD camera (Photometrics), cooled to −70 °C and operated through MicroManager (42) with 33 ms exposure at 200 electron multiplier gain. Pixel width in the magnified image was 96 nm, determined using a USAF calibration target (Edmund Optics). An excess of TerL during slide preparation was required to observe multi-step photobleaching events, though this led to a higher background of spots that photobleached in one step. Despite an excess of TerL, the proportion of DNA co-localising with protein was low. The packaging assembly was therefore likely unstable at the concentrations used. Colocalization was determined based on whether the intensity-weighted centroid of the GFP signal is within experimental error of the BOBO-3 signal centroid. We define experimental error as the precision with which the x- and y-position of one GFP monomer could be assigned in time relative to another monomer as described previously (43) (see Supplementary Note). For each event, the GFP fluorescence intensity trace was subjected to Chung-Kennedy filtering (44) and step-wise photobleaching was fitted using the Progressive Idealization and Filtering (PIF) algorithm (45). Over-fitted events were rejected according to the chi-squared statistic-based criterion as implemented for the analysis of microtubule assembly data (46).

### Cryo-electron microscopy imaging and reconstruction

DNA packaging reactions containing 125 nM prohead, 125 nM *Sca*I-linearised pUC18 DNA, TerL and TerS at 1 μM and 2 μM monomer concentrations, respectively, were applied to Quantifoil R 2/2 copper grids (Quantifoil Micro Tools GmbH, Jena, Germany) within ∼30 s after addition of ATP to 1 mM final concentration, then blotted and vitrified using a Vitrobot Mark III. Data were collected on a Tecnai Polara cryo-electron microscope operating in nanoprobe mode at 300 kV accelerating voltage and at a nominal magnification of 78000×. Using EPU, images were recorded on a Falcon 2 detector with an image pixel size of 1.37 Å/px. 30 frames were collected per micrograph with a dose rate of 40 e^-^/Å^2^/s and total exposure time of 1.65 s. All cryo-EM equipment and control software were supplied by Thermo Fisher Scientific, Waltham, MA. Drift correction was performed using MotionCor2 (47), and CTF estimation using CTFFIND4 (48). Prohead particles were picked automatically (icosahedral reconstruction) or manually (asymmetric reconstruction) in X3D (49). All reconstructions were performed with AUTO3DEM according to gold standard (50). For asymmetric reconstruction, the manually oriented particles were first subject to an icosahedral reconstruction procedure to determine accurate particle centers. Angles were then reassigned to approximate values defined by the portal vertex recorded during manual picking, and a local angular refinement was performed in AUTO3DEM around these values with a regular icosahedral capsid as initial model.

### Native mass spectrometry analysis

High-resolution native MS measurement was performed using a modified Q-Exactive hybrid quadrupole-Orbitrap mass spectrometer (Thermo Fisher Scientific) optimized for high mass measurement and retention of non-covalent interactions (51, 52). TerS protein was buffer-exchanged into 200 mM ammonium acetate, pH 7.5 using Micro Bio-Spin P-6 Gel columns (Bio-Rad) at 4 °C. Ions were generated in the positive ion mode from a static nanospray source using gold-coated capillaries prepared in house. The instrument was operated in “native mode” with a wide-pass isolation window. Transmission parameters used were as follows: capillary voltage 1.2kV, capillary temperature 80°C, inject flatapole 7 V, inter flatapole lens 6 V, transfer multipole 4 V, C-trap entrance lens 5.8 V. Ions were trapped in the higher-energy collisional dissociation (HCD) cell before being transferred into the C-trap and Orbitrap mass analyzer for detection, with 100 V applied to the HCD cell to aid ion de-solvation. Argon was used as the collision gas and the pressure in the HCD cell was increased until UHV pressure was ∼1.2 × 10^−9^ mbar. Transient time was 64 ms (resolution of 17,000 at m/z 200), automatic gain control (AGC) target was 1 × 10^6^ with a maximum fill time of 100 ms, micro-scans were set to 15, with no averaging, and the threshold parameter was set to 3 (reduced from default of 4.64). Calibration using clusters of CsI was performed prior to mass measurement up to m/z 11,304. Spectra were averaged using Xcalibur 2.2 (Thermo Fisher Scientific) and deconvoluted using Unidec 2.7.3 (53). To estimate the error in the fit, the mass was calculated from the four most abundant charge states using the MaCSED software tool available from http://benesch.chem.ox.ac.uk/resources.html.

## RESULTS

### Reconstitution of a HK97 DNA packaging system

To reconstitute an *in vitro* DNA packaging system based on bacteriophage HK97, we combined recombinant HK97 TerS and TerL proteins with empty, portal-containing procapsids isolated from *Escherichia coli* infected with a previously characterised packaging-deficient mutant amC2 (31). We showed by DNase protection assays that the assembled system can package viral genomic DNA isolated from wild-type bacteriophage into procapsids (Figure 1A and B). Intriguingly, the assembled system could also package linearised plasmid DNA, irrespective of whether the HK97 *cos* site was present, and irrespective of whether the DNA substrate had blunt ends or 5′ or 3′ overhangs (Figure 1C). Furthermore, the presence of TerS appeared to stimulate DNA packaging (Figure 1D).

**Figure 1.**
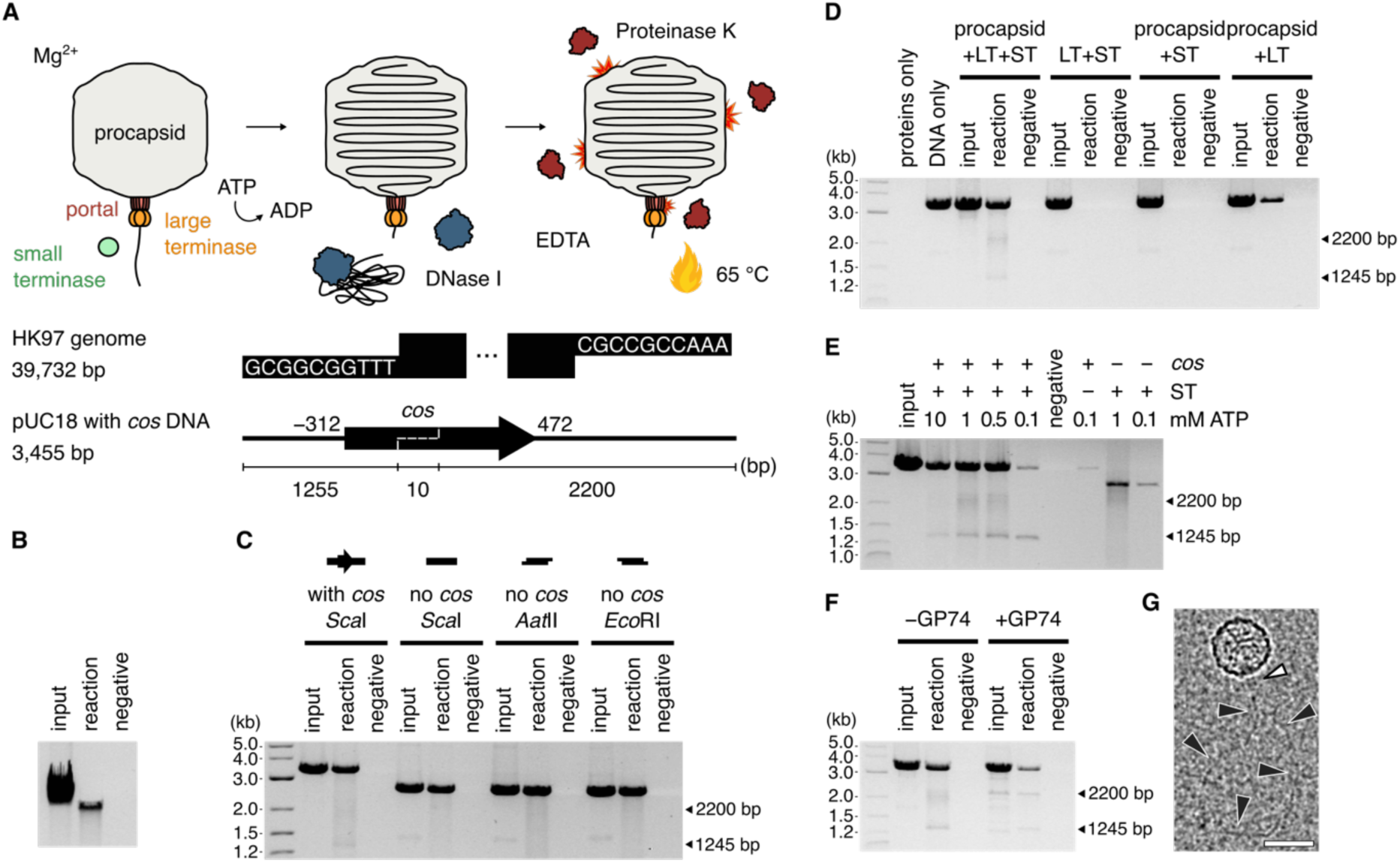
Reconstituting the HK97 DNA packaging system. (**A**) Schematic of the DNA packaging assay and DNA substrates. The HK97 genome boundary is defined by *cos* ends which contain 10-nt 3′-overhangs. A 784-bp region around the *cos* end (−312..+472, dashed line) was inserted into pUC18 plasmid DNA and the DNA linearized with *Sca*I. Under conditions of excess TerL and TerS proteins, ATP was added to initiate packaging of DNA through the portal complex of purified procapsids in solution. (**B**) Packaging of the HK97 genome. Input, reaction with no DNase treatment. Reaction, with DNase treatment. Negative, reaction with no ATP but with DNase treatment. (**C**) Packaging of linear pUC18 DNA with or without *cos* DNA, with blunt (*Sca*I) or sticky ends (*Aat*II, *Eco*RI). (**D**) Packaging of *cos*-containing DNA using different protein combinations (LT, large terminase; ST, small terminase). (**E**) ATP titration and stalling of the HK97 motor. (**F**) GP74-induced cleavage of the DNA substrate. Arrows in (C–F) indicate the two non-full-length fragments protected and the expected size of products after cleavage at the internal *cos* site of the linearized DNA substrate. (**G**) Cryo-EM micrograph of the reconstituted DNA packaging assembly. Scale bar = 50 nm. White arrowhead indicates a terminase assembly on the portal of the HK97 procapsid. Black arrowheads indicate the path of DNA emerging from the terminase assembly.

The ability of the system to package any short linear substrate suggested that DNA translocation can be uncoupled from *cos* site-specific initiation *in vitro* as long as a free DNA end is provided. Making use of this property, we further probed the biochemical requirements of efficient DNA translocation using linearised plasmid DNA (Figure S1). We found that packaging required magnesium or manganese and was inhibited by zinc. The system was active from pH 6 to 9 and tolerated up to 300 mM potassium glutamate or 150 mM sodium chloride. The charge-neutralising spermidine was essential for the packaging of long genomic DNA (39.7 kb) but dispensable for the packaging of shorter linear DNA (3.5 kb). These observations indicated that the HK97 system responds to pH, divalent metals and monovalent salts similarly to other DNA packaging systems such as lambda (54) and phi29 (19), and even other NTP-dependent DNA-processing machineries such as DNA polymerase III (55).

In addition to protection of the full substrate, we observed protection of shorter DNA fragments when an intact internal *cos* site was present (Figure 1C to E). The sizes of these fragments corresponded to the cleavage products of a *cos* cleavage reaction, which are 1245 bp and 2200 bp in length. However, the absence of fragments in the non-DNase-treated controls (input) indicated there has been no cleavage of the substrate; rather, the motor had stalled on the *cos* site at the point when DNase was added. This behavior was dependent on the presence of TerS protein and became less apparent with increasing ATP concentrations (Figure 1E). We reasoned that an internal *cos* site in the substrate would represent a downstream *cos* signal at the next boundary between genome copies in the viral DNA concatemer. TerS bound to the *cos* site could thus present a roadblock to the translocation machinery, while an excess of ATP helped by ensuring continual turnover by TerL, increasing its chances of overcoming this roadblock. We note that the ∼1.3 kb fragment was more defined and stronger in intensity at all ATP concentrations tested, indicating that the roadblock was more specific and efficient in the forward direction, which perhaps confers a directionality in this packaging system.

HNH endonuclease GP74 has been reported to stimulate *cos* DNA cleavage *in vitro* by TerL (32). Indeed, addition of recombinantly produced GP74 to the packaging reaction caused the above observed DNA fragments to appear in the non-DNase-treated controls (input), signifying cleavage (Figure 1F), with the ∼2.2 kb fragment also appearing more defined. Taken together, our biochemical observations suggest that the minimal components for DNA packaging *in vitro*, when the substrate is linear and has free ends, are TerL and the HK97 procapsid only. TerS is not essential, but stimulates packaging and contributes to stalling and possibly site-specific cleavage at a downstream *cos* signal, perhaps acting in tandem with the slowing of translocation as observed in other packaging systems. Excitingly, cryo-EM imaging of the assembled system at a sparse particle density revealed protrusions from the procapsid at the unique portal vertex followed by DNA-like densities (Figure 1G), which suggested the HK97 system assembles with a similar architecture as the *pac* and phi29 systems.

### Structure of the HK97 TerL reveals a classical viral packaging ATPase and nuclease

To gain structural insights into the active ATPase of the HK97 system, we determined the crystal structure of the *apo* HK97 TerL at 2.2 Å resolution (Figure 2A and Table S1). The structure comprises both ATPase and nuclease domains, with the two connected by a short linker and a unique β-strand (Figure 2A and B, red) that is absent in *pac* TerL proteins. This unique β-strand forms a three-stranded β-sheet with the auxiliary hairpin of the nuclease domain (Figure 2A and B, beige). An additional ααβ element was observed within the nuclease domain relative to *pac* TerL proteins (Figure 2 and Figure S2, blue), possibly serving as a platform for interactions with HNH endonuclease GP74 or simply part of an adaptation of *cos* TerL proteins towards sequence-specific endonuclease activity.

**Figure 2.**
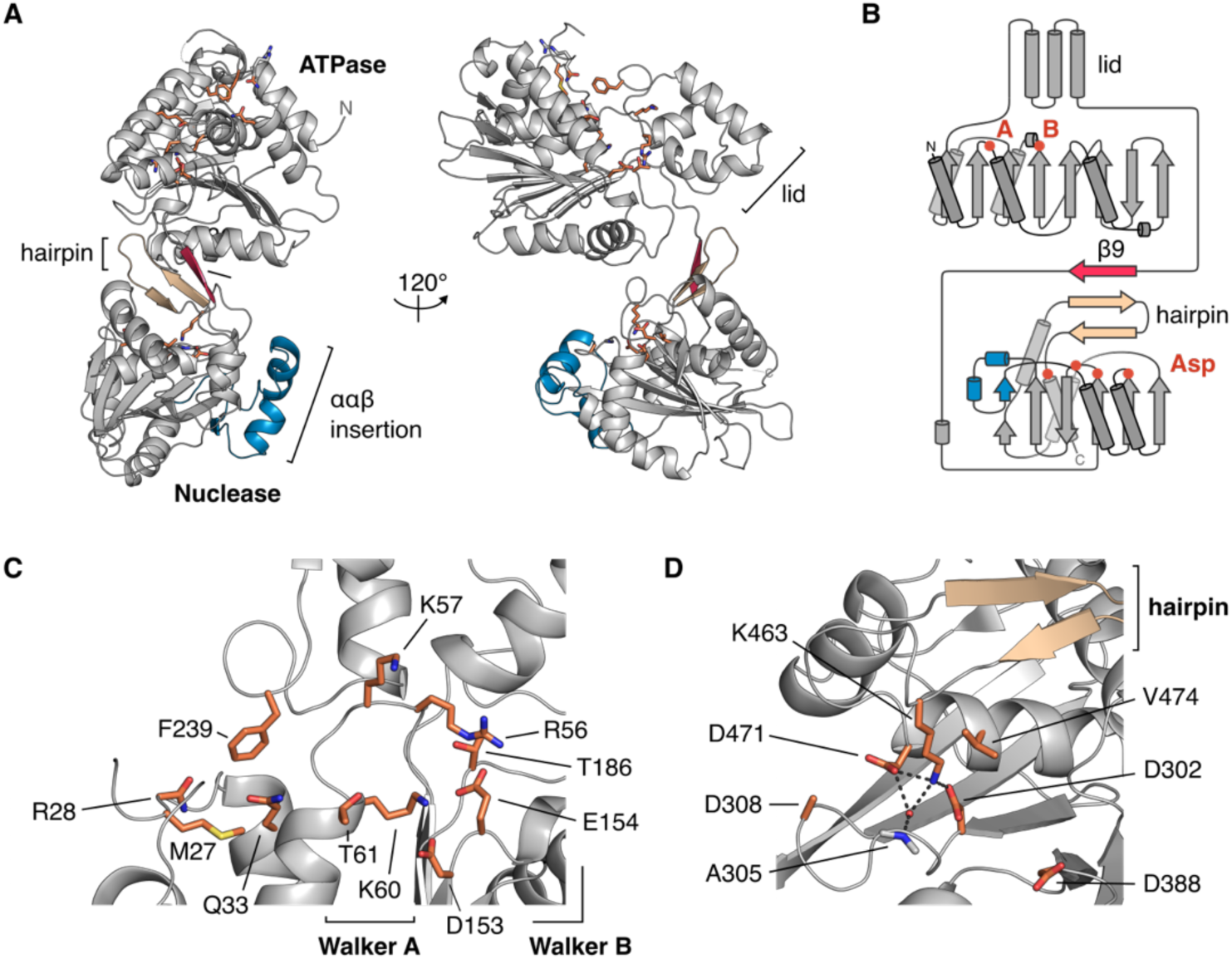
Crystal structure of the HK97 TerL. (**A**) Ribbon representation. Orange sticks depict the putative ATPase and nuclease active sites. Secondary structure elements unique to the HK97 protein are colored in red and blue. The conserved β-hairpin of viral terminase nucleases is colored in beige. Topology diagram of the HK97 protein. The positions of Walker A, B residues, and aspartate residues in the nuclease active site are marked. View of the (**C**) ATPase and (**D**) nuclease catalytic centers.

Movement between the ATPase core and lid during ATP turnover has been implicated in driving DNA translocation. The ATPase core comprises an eight-stranded β-sheet, with six parallel strands and two anti-parallel strands (Figure 2), a topology that is conserved across viral packaging ATPases, which build on the ASCE 51432 topology (Figure S3A) (56). The Walker A motif is found between strands β1 and β2. Unlike in lambda gpA, a lysine residue, K60, occurs at the expected position; however, like in lambda (57, 58), a second lysine residue, K57, occurs N-terminal to the classical position (Figure 2C and Figure S3B). Interestingly, this second lysine residue is conserved within herpes virus TerL proteins (Figure S3B). The Walker B motif resides at the end of strand β6 with catalytic residues D153 and E154 (Figure 2C). Further downstream is residue T186, which superposes well with the C-motif of the T4 TerL gp17 or, in general, the sensor I residue of ASCE ATPases, a hydrogen-bonding residue that helps to position water molecules for nucleophilic attack of ATP (59). A putative adenosine-binding pocket is formed by a conserved glutamine residue (3), Q33, and hydrophobic residues F239 and M27 (Figure 2C).

The HK97 TerL nuclease domain has an RNase H-like fold, with residues D302, D388 and D471 at the catalytic center. Adjacent to the catalytic center, we found an auxiliary β-hairpin that is conserved among *pac* TerL nuclease domains (Figure 2D). Residue K463, which follows the hairpin, protrudes into the active site, forming potential salt bridges with D302 and D471 and coordinating a water molecule. A similarly placed lysine in *pac* phage Sf6 has been proposed to act as a switch for metal binding (60) and alters DNA accessibility to the nuclease domain (61). As stated earlier, a β-strand stacks against the hairpin and connects the nuclease domain with the ATPase lid. We speculate that this structure could play a role in switching TerL activity between DNA cleavage and translocation. Despite local variations in the Walker A sequence and secondary structure elements in both ATPase and nuclease domains, the HK97 TerL shows high structural homology to its *pac* counterparts, hinting at a common set of structural mechanisms that regulate their activity.

### The HK97 TerS forms a circular 9-subunit assembly

Our biochemical data indicated that the HK97 TerS can stimulate and arrest DNA packaging. However, the structure and stoichiometry of an *apo* cos TerS have remained elusive. We addressed this by in-solution biophysical methods and X-ray crystallography. By size exclusion chromatography with multi-angle laser-light scattering (SEC-MALS) and native mass spectrometry, the HK97 TerS measured 160.5 kDa and 165.020 ± 0.005 kDa, respectively, in mass (Figure S4), corresponding to an oligomer of nine 18.3-kDa subunits. Additionally, we determined the crystal structure of the HK97 TerS to 1.4 Å resolution (Figure 3A and Table S2). The protein crystallized with three subunits in the asymmetric unit around a crystallographic 3-fold axis, forming ultimately a 9-subunit circular oligomer.

**Figure 3.**
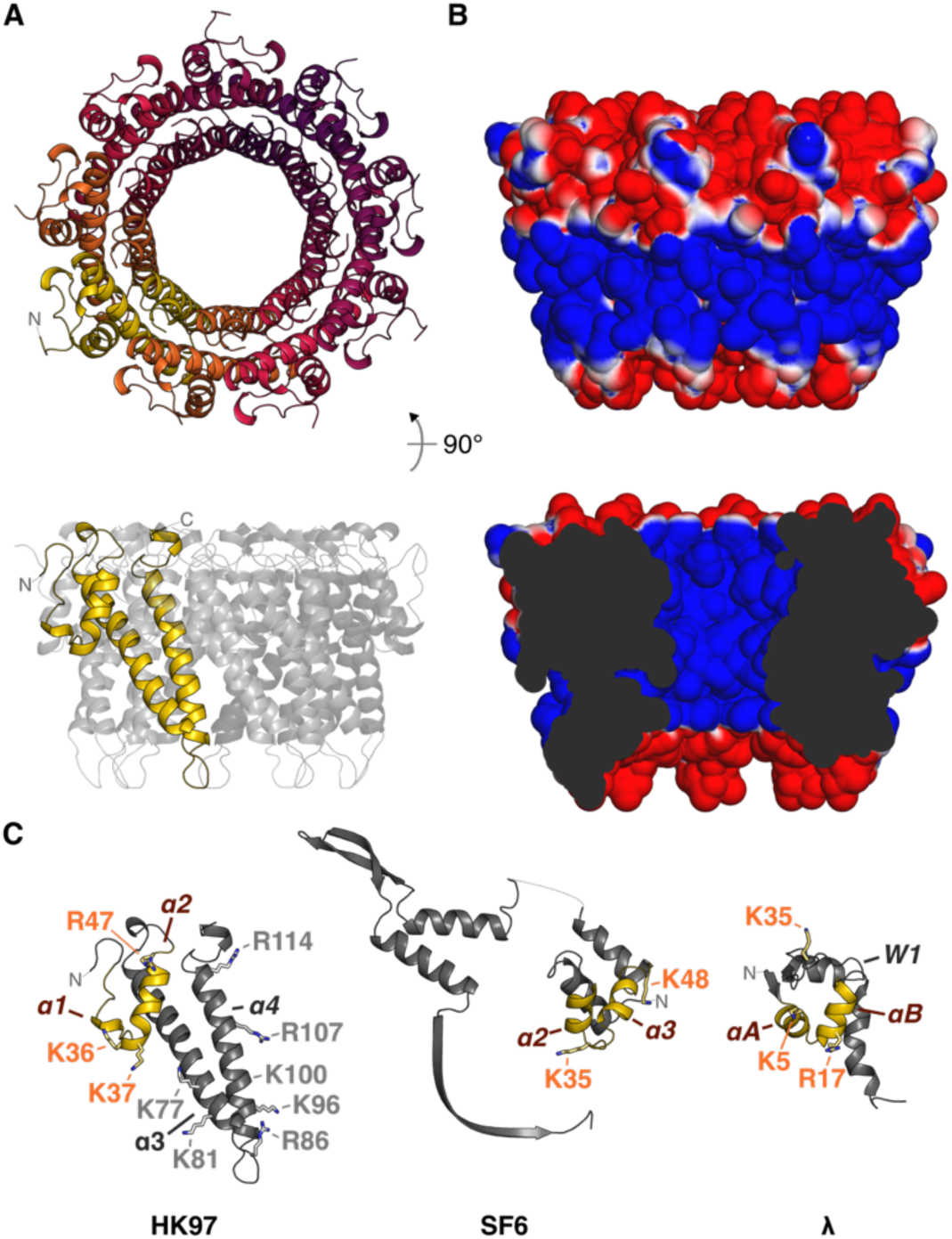
Crystal structure of the HK97 TerS. (**A**) Ribbon representation of the TerS oligomer. (**B**) Solvent accessible surface of the TerS oligomer, colored by electrostatic potential from −1 to 1 kT/*e*. Comparison of subunit structure with *pac* phage SF6 GP1 (PDB 3ZQQ) and *cos* phage lambda gpNu1 (PDB 1J9I). Putative DNA-binding helix-turn-helix motifs are colored in yellow. Residues important for DNA binding in SF6 and lambda are displayed as sticks. Corresponding residues in the HK97 TerS are indicated with orange text.

Of the 161 residues in each protein chain, residues 21–124 had well-resolved electron density. The N-terminal region superposes loosely with the a2 and a3 helices of the helix-turn-helix motif of *pac* phage SF6 GP1 (62) and of the winged helix-turn-helix domain of lambda gpNu1 (63) (Figure 3C). Two strong helix-breaking proline residues, P28 and P29, occur N-terminally and prevent an a1 helix from forming. Despite this, an overall HTH-like structure is maintained owing to a local network of intramolecular interactions (Table S3). Following the N-terminal region, two long helices form the basis of oligomerization for the HK97 TerS oligomer (Figure 3C). The HK97 oligomer has a van der Waals outer diameter of roughly 85 Å, and an inner tunnel 55 Å long and 18 Å wide at its narrowest point. Residues K36, K37, R47, K77 and K81 form a positive belt around the outside of the complex (Figure 3B). However, positive charges from R86, K96, K100, R107 and R114 also line the inner tunnel, which is solvent-exposed and almost wide enough to accommodate double-stranded DNA. Two models of DNA binding have been proposed for TerS proteins in general, with DNA wrapped around the oligomer or feeding through the oligomer. Our structure presented here does not confirm or reject either of these possibilities. At the C-terminus, 37 residues are unmodelled. These could assist with DNA binding as has been shown in *pac* phage P22 (17). Though the exact mechanism of DNA binding remains unknown for *cos* TerS proteins, similarities in structure and stoichiometry suggest that HK97 and *pac* viruses employ a similar viral DNA recognition mechanism to initiate packaging, but it remains to be seen how specificity is conferred for packaging arrest and termination.

### The HK97 TerL assembles into a pentamer for DNA translocation

TerL proteins of *pac* bacteriophages pentamerize when they assemble to form an active motor (3). To ascertain the stoichiometry of the HK97 TerL in an active DNA-translocating complex, we conducted a single-molecule subunit counting assay, where we monitored the step-wise photobleaching of N-terminal GFP-TerL fusion proteins by total internal reflection fluorescence microscopy (Figure 4A). We initiated packaging with a 230-bp DNA fragment that encompassed the HK97 *cos* site and was 5′-biotinylated on the downstream end, in the presence of TerS and ATP. Stalling immediately with ATP-γ-S, we tethered the complex to a streptavidin-coated slide surface for imaging. We detected spatially co-incident GFP-TerL and DNA-staining BOBO-3 signal to localise individual complexes on the imaging surface. The number of GFP photobleaching steps under continuous illumination indicated the number of TerL subunits in a stalled complex (Figure 4B). We counted a range of GFP photobleaching steps coincident with DNA (Figure 4C, Figure S5, and Table S4). Accounting for a naturally occurring non-fluorescent GFP population (45), missed steps due to fast consecutive photobleaching, and error in step detection (see Supplementary Note), we conclude that the HK97 DNA packaging assembly has an upper limit of 5 TerL subunits.

**Figure 4.**
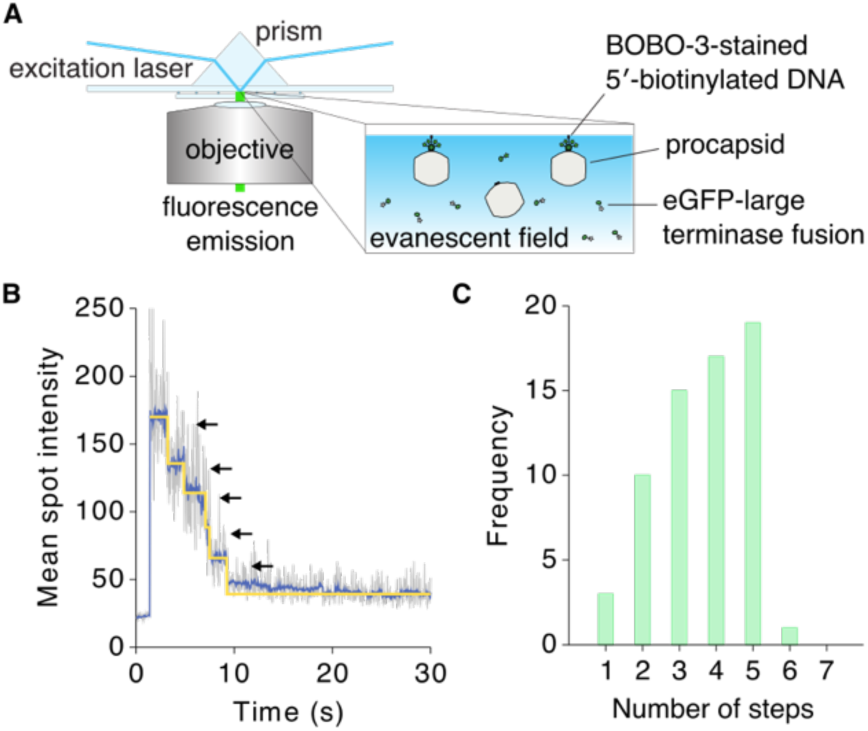
Single-molecule photobleaching analysis of large terminase stoichiometry. (**A**) BOBO-3-stained 5′-biotinylated DNA was pre-incubated with procapsids, TerS and GFP-labelled TerL protein. ATP was added to initiate translocation and chased with ATP-γ-S. The complex was immobilized on a streptavidin-coated surface for visualization by total internal reflection fluorescence microscopy. GFP, λ_ex_/λ_em_ = 488 nm/509 nm. BOBO-3, λ_ex_/λ_em_ = 570 nm/602 nm. (**B**) GFP fluorescence trace of one protein spot that co-localized with DNA. (**C**) Number of photobleaching steps observed for protein spots co-localizing with DNA (*n*=65), which indicated an upper limit of 5 TerL subunits.

In the aforementioned cryo-EM imaging of the packaging complex at sparse particle density, we obtained 1024 particles where we could define the long axis of the particle based on portal density. Using the protocol described in the Material and Methods, where icosahedral and 5-fold symmetry was applied, we were able to unambiguously resolve the procapsid in association with TerL at the unique portal-containing vertex of the procapsid (Figure S6A and B). Six-fold averaging around the long axis revealed a dodecameric portal that extends ∼140 Å inside the procapsid and contains all characteristic domains (64), including the crown, wing, stem and the clip (Figure S6C, inset). Disordered densities distal to the portal suggested a dynamic terminase/DNA assembly. We imposed 5-fold symmetry and docked into this distal density a HK97 TerL model generated based on our crystal structure and the *pac* phage P74-26 model, to create a hybrid model of the HK97 DNA packaging assembly. This superposition did not reveal a clear candidate for a *trans*-acting arginine finger, but provided a working model where the ATPase lid of one HK97 TerL subunit inserts between the ATPase core and nuclease domain of an adjacent subunit to serve as an anchor point for DNA translocation, as proposed for P74-26 (2).

### An unusual critical lysine residue for HK97 TerL ATPase activity and DNA translocation

In solution, the HK97 TerL appeared as a monomer, measuring 52.7 kDa and 52.5–56.7 kDa in mass by SEC-MALLS and AUC, respectively (Figure S7). The protein was a poor ATPase as a monomer at physiological ATP concentrations, but on formation of a packaging complex, it became an active ATPase (Figure 5A and B). Since this has direct implications on ATP usage during virus assembly in the cell, we wanted to understand what drives and regulates ATPase activity in the HK97 TerL.

**Figure 5.**
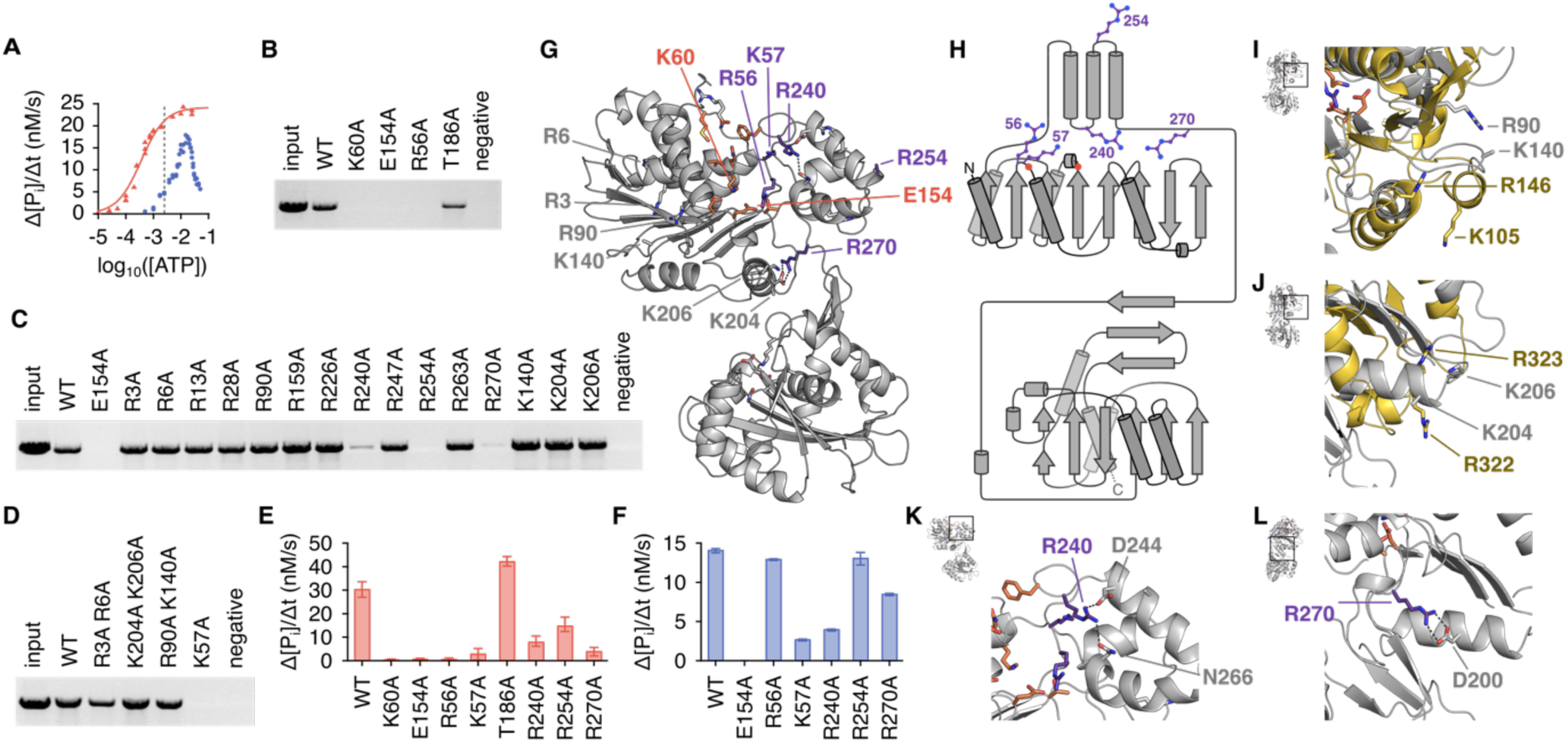
Critical residues for HK97 TerL ATPase activity and DNA translocation. (**A**) ATP hydrolysis rates, based on inorganic phosphate release, of free TerL (blue) and TerL in the presence of DNA, TerS and procapsids (red) as a function of ATP concentration. Fitting of the Hill equation suggested a *V*_max_ of 22.7 nM/s, *K*_m_ of 0.32 mM and *n* of 1.4 for TerL in the DNA packaging assembly. The average physiological ATP concentration, 1.5 mM (73), is indicated with a dotted line. (**B** to **D**) DNA packaging by ATPase alanine TerL mutants. (**E**) ATPase activity of mutant TerL in the presence of DNA, TerS and procapsids at 1 mM ATP concentration. (**F**) ATPase activity of free mutant TerL at 10 mM ATP concentration. (**G**) Ribbon representation of the HK97 TerL structure with ATPase active site residues indicated in orange, and other catalytically important arginine or lysine residues purple. (**H**) Topology diagram with the same residues marked. Superposition of the HK97 structure with (**I**) the *trans*-acting residues of the phi29 packaging ATPase (gold, PDB 7JQQ) and (**J**) the arginine finger of NTP-binding domain 1 of *Thermus thermophilus* AAA+ ATPase ClpB (gold, PDB 1QVR). (**K, L**) Side-chain interactions involving residues R240 and R270, respectively.

Mutation of Walker A residue K60 and Walker B residue E154 to alanine abrogated DNA packaging and ATPase activity as expected (Figure 5B, E and F). However, mutation of putative Sensor I residue T186 had little effect, suggesting that the residue does not act as a Sensor I as proposed for T4 gp17. Mutation of R56 affected function. R56 corresponds to R162 of T4 gp17, thought to be a *cis*-acting arginine finger that repositions upon oligomerization to allow contact with the ATP γ-phosphate (65). The same residue in P74-26 TerL, R39, was thought to be a sensor II-like residue, which instead of having a catalytic role, coordinates conformational changes during ATP turnover for mechanical movement. More recent work on lambda gpA suggested that this arginine toggles between the catalytic Walker B glutamate and a glutamate residue in the ATPase lid to coordinate movement during ATP turnover (26). However, there is no glutamate residue in the HK97 ATPase lid that approaches R56 in our *apo* structure. Therefore, the role played by R56 in DNA translocation remains ambiguous. Nevertheless, it remains an important and conserved arginine residue for viral packaging ATPases.

*Trans*-acting elements important for nucleotide exchange or catalysis have been identified in P74-26 and phi29 on the opposite side of the β-sheet away from the catalytic center (1, 2) (Figure S3A). While no positive residues occur in the corresponding vicinity of phi29 residue R146, a residue known to be important for nucleotide exchange, K140 of the HK97 TerL aligns well with the arginine finger of both phi29 and P74-26. However, mutation of K140 to alanine did not impair DNA packaging (Figure 5C and I, and Figure S8). Mutation of R90 in the vicinity also did not impair packaging. Furthermore, mutation of K204 and K206, which align with the double fingers of *Thermus thermophilus* AAA+ATPase ClpB (66), did not impair activity (Figure 5C, D and J). These results suggest that there are no classical arginine fingers in the HK97 TerL. To widen our search, we generated alanine mutants for every arginine residue in the domain and every lysine residue on the surface of the β-sheet distal to the catalytic center (Figure 5G and H). In case of redundancy in closely spaced residues, double mutants were also generated. Of all mutants tested, only R240A, R254A and R270A had reduced activities (Figure 5C, E and F), which were not due to protein unfolding (Figure S8A). These mutants could not be rescued by addition of the Walker B E154A mutant (Figure S8C and D), which supplemented the system with intact arginine fingers, indicating that the residues do not participate in catalysis in *trans*. Given the local environment, we speculate that R240 and R270 mediate movement between the ATPase core and lid through salt bridges and hydrogen bond interactions during ATP turnover (Figure 5K and L), and though not essential for hydrolysis, R254 may contact DNA or have a role in mechanochemical coupling during translocation of the DNA.

Contrary to P74-26 TerL and the phi29 ATPase, the HK97 TerL ATPase contains a second lysine residue upstream of the Walker A lysine. Unexpectedly, mutation of this residue, K57, abrogated DNA packaging and ATPase activities (Figure 5D, E and F). In the absence of a *trans*-acting arginine finger, K57 could provide the additional charge needed for ATP coordination. As mentioned, herpes viruses and phage T5 also encode an additional lysine upstream of the Walker A lysine (Figure S3B). Thus, a second lysine at this position may be a defining feature of a previously uncharacterised subfamily of viral packaging ATPases, with subtly different ATPase regulatory mechanisms. Given the differences in activity observed between free TerL and TerL during DNA translocation, however, a conformational switch likely still exists to activate the ATPase domain upon oligomerization.

## DISCUSSION

Studies on bacteriophage HK97 have enabled several important discoveries, in particular, the identification of the universal HK97 capsid protein fold (67), which is now known to be conserved among tailed bacteriophages and herpes viruses (68, 69). Just as they share a capsid protein fold, many phages and herpes viruses encode similar packaging machinery for the insertion of their genomes into capsids (70). However, mechanistic differences exist between the packaging machineries of different viruses, stemming from a difference in terminase structure and in the overall architecture of the active packaging motor (1, 3, 71). The outcome is a packaged genome with different but characteristic ends depending on the packaging strategy used: *cos, pac* or phi29-like.

The packaging machineries of *pac* and phi29-like systems are well characterised. However, the lack of molecular structures for *cos* TerL and TerS proteins such as those of lambda and HK97 has limited our understanding of the evolutionary relationships among these viruses in the context of packaging. In our reconstitution of the HK97 packaging assembly, we discovered an important trait that marks HK97 as a *cos* virus, that is, the ability of TerS to facilitate sequence-specific stalling of the motor at an intact *cos* DNA signal (Figure 1E). However, our study also revealed unexpectedly strong similarities between the HK97 terminase proteins and their *pac* counterparts, both in structure and in biophysical properties. The HK97 TerS has similar N-terminal HTH-like elements and exists as a stable, circular 9-mer assembly in solution (Figure 3 and Figure S4). Likewise, the ATPase and nuclease domains of the HK97 TerL are similar to their *pac* counterparts (Figure 2, Figure S2 and Figure S3). Our photobleaching-based analysis suggested that the HK97 TerL, too, oligomerizes on the portal of the procapsid to form a pentamer (Figure 4). Lambda gpNu1 and λ has been found to associate in a 2:1 ratio in solution, and tetramerize to form the maturation complex for DNA processing (9, 71). We speculate that a substantial rearrangement in TerL could occur in lambda to form the DNA translocation complex. Equally, the difference in TerL stoichiometry may also be an inherent difference between HK97 and lambda systems. As such, our results would place HK97 in a subset of *cos* phages that are evolutionarily closer in DNA packaging mechanisms to *pac* and phi29 viruses.

At cellular ATP concentrations, the HK97 TerL ATPase becomes activated only upon formation of the packaging assembly (Figure 5). Despite a complete mutagenesis study of arginine residues in the ATPase domain, we were unable to identify a classical *trans*-acting arginine finger that could be rescued by a Walker B active site mutant. This is unlike in *pac* phage P74-26 and in phi29 (1, 2, 22). Instead, the HK97 ATPase contains an additional critical positive residue, K57, in the Walker A region. This residue, along with R56, could provide the electrostatic potential required for ATP discrimination and coordination of ATP hydrolysis in the absence an arginine residue from a neighbouring subunit. An alternate trans-regulatory mechanism has been proposed recently for P-loop NTPases, which involves monovalent cation binding stabilising ATP in a catalytically competent conformation (72). The exact conformational switch that activates HK97 TerL upon oligomerization is still unknown. However, we identified additional residues around the ATPase lid that contribute to DNA packaging with possible roles in mechanochemical coupling or inter-subunit communication.

Cleavage of *cos* DNA in lambda generates 10-nt 5′ overhangs, whereas in HK97 12-nt 3′ overhangs are generated (27). Our *in vitro* packaging system did not distinguish between DNA ends. Thus, DNA translocation and packaging initiation are seemingly uncoupled. Unlike lambda, HK97 encodes an HNH endonuclease, GP74 (32). As observed before and reproduced in our *in vitro* system, GP74 promotes cleavage at the *cos* site (Figure 1F). Compared to homologues among viruses, the HK97 TerL nuclease features additional secondary structure elements (Figure 2). Since the structure of lambda gpA is still unknown, it is yet unclear if these elements are a general feature of *cos* TerL proteins for site-specific DNA cleavage or regulation between the ATPase and nuclease domains, or a specific feature for protein-protein interactions in viruses that encode an HNH endonuclease.

This study establishes the minimal components required for *in vitro* packaging by *cos* bacteriophage HK97, a widely studied viral model system. This breakthrough has enabled us to address long-standing questions about the biochemical and structural differences between a *cos* packaging system and other well-characterized archetypal packaging systems (*pac* and phi29) employed by double-stranded DNA viruses. We have presented here the first atomic structures of a TerL and TerS protein from a *cos* bacteriophage, and with them stoichiometric information on TerS in isolation and on TerL as part of an *in vitro* reconstituted motor. Our biochemical, structural and biophysical analyses highlight that there are no specific criteria based on overall protein fold that delineate a *cos* phage, a *pac* phage and a phi29-like phage; rather, a continuum of assembly and regulatory strategies likely exists and this has been adapted in the HK97 phage to package unit-length genomes into capsids.

## ACCESSION NUMBERS

Atomic coordinates and structure factors for the reported TerL and TerS crystal structures have been deposited with the Protein Data bank under accession numbers 6Z6D and 6Z6E, respectively. Cryo-EM density maps of the DNA packaging assembly are deposited with the Electron Microscopy Data Bank under accession numbers EMD-22099, EMD-22100 and EMD-22101.

## ACKNOWLEDGEMENT

The authors would like to thank A. Leech for support and access to the University of York Bioscience Technology Facility, J. Turkenburg and S. Hart for support in crystallographic data acquisition, the York Advanced Research Computing Cluster, and the Diamond Light Source. The expression plasmid encoding GP74 was a gift from K. L. Maxwell, University of Toronto, Canada. Two of the authors (Dr. Shelley Grimes and Prof. Roger Hendrix), who made important contributions at the early stages of this work, are now deceased. The other authors would like to dedicate this work to their memory.

