## Supplementary Material for "Structural basis of DNA packaging by a ring-type ATPase from an archetypal viral system"

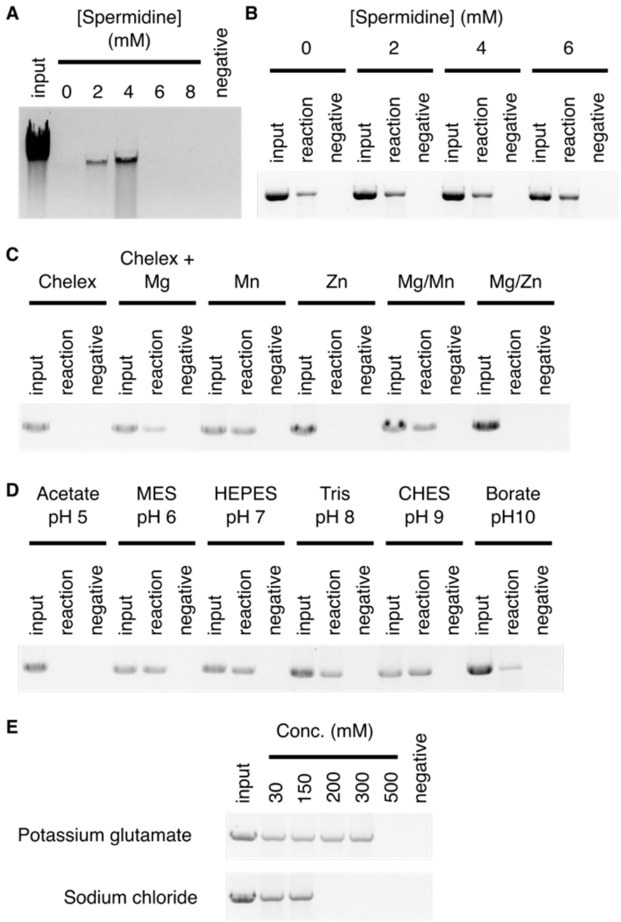


**Figure S1. Biochemical properties of the HK97 DNA packaging system.** Dependence on spermidine in the packaging of (**A**) the HK97 genome and (**B**) linear *cos*-containing pUC18 DNA. Dependence on (**C**) divalent metal ions, (**D**) pH and (**E**) monovalent salts in the packaging of linear cos-containing pUC18 DNA. Reaction components were treated with Chelex-100 resin before addition of divalent metal ions. Samples with different salt levels were ethanol-precipitated before gel analysis.

**Table S1. TerL crystallographic data collection and refinement statistics.**

| | SeMet | | KBr soak |
|---|---|---|---|
| | *peak* | *inflection* | |
| **Data collection** | | | |
| Wavelength (Å) | 0.97949 | 0.97969 | 0.91983 |
| Space group | *C*2 | *C*2 | *C*2 |
| Cell dimensions | | | |
| *a*, *b*, *c* (Å) | 211.5, 39.3, 66.0 | 211.9, 39.3, 66.1 | 211.8, 39.2, 65.5 |
| $\beta$ (°) | 103.3 | 103.2 | 103.4 |
| Resolution (Å) | 49.59–2.00 (2.05–2.00) | 49.72–2.40 (2.49–2.40) | 45.53–2.20 (2.27–2.20) |
| $R_{merge}$ | 0.122(1.233) | 0.100 (1.249) | 0.136 (0.842) |
| $I / \sigma(I)$ | 10.1 (1.0) | 13.6 (1.6) | 7.1 (1.8) |
| $CC_{1/2}$ (%) | 99.7 (36.5) | 99.9 (52.9) | 99.1 (59.5) |
| Completeness (%) | 98.3 (85.3) | 99.9 (99.7) | 99.6 (98.2) |
| Redundancy | 6.2 (4.1) | 6.6 (6.4) | 3.3 (3.3) |
| Wilson B (Å$^2$) | 43.3 | 60.5 | 37.6 |
| No. unique reflections | 35605 (2257) | 21206 (2198) | 26796 (2242) |
| **Refinement** | | | |
| Resolution (Å) | | | 45.53–2.20 |
| No. reflections | | | |
| Working | | | 25551 |
| Free | | | 1244 |
| $R_{work}/R_{free}$ | | | 0.205/0.254 |
| No. atoms | | | |
| Protein | | | 3522 |
| Water | | | 93 |
| Ligand | | | 18 |
| B factors | | | |
| Protein | | | 38.9 |
| Water | | | 33.3 |
| Ligand | | | 53.1 |
| R.m.s. deviations | | | |
| Bond lengths (Å) | | | 0.008 |
| Bond angles (º) | | | 1.3 |
| Ramachandran | | | |
| Favored (%) | | | 97.5 |
| Outlier (%) | | | 0.0 |

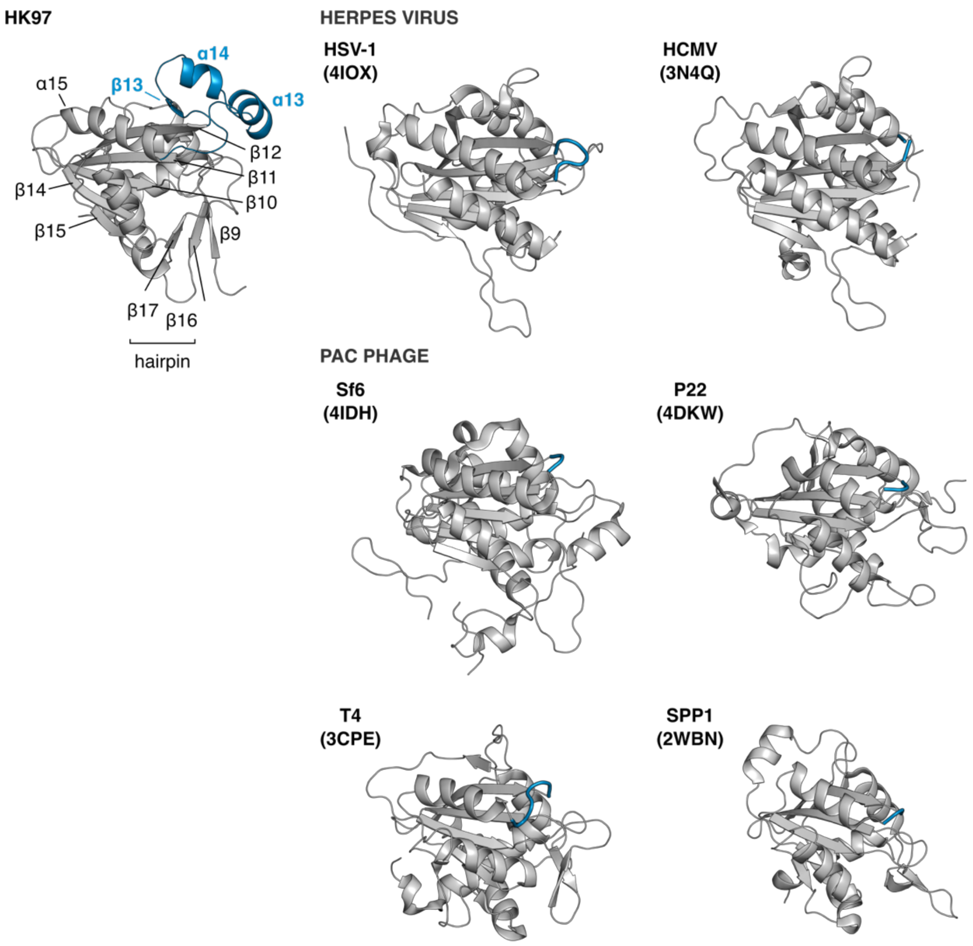


**Figure S2. Comparison of viral packaging nucleases.** The HK97 TerL nuclease contains an additional ααβ element, colored blue, on the periphery of the central β-sheet. The corresponding region in herpes virus and *pac* homologues is colored blue. PDB codes are indicated in brackets.

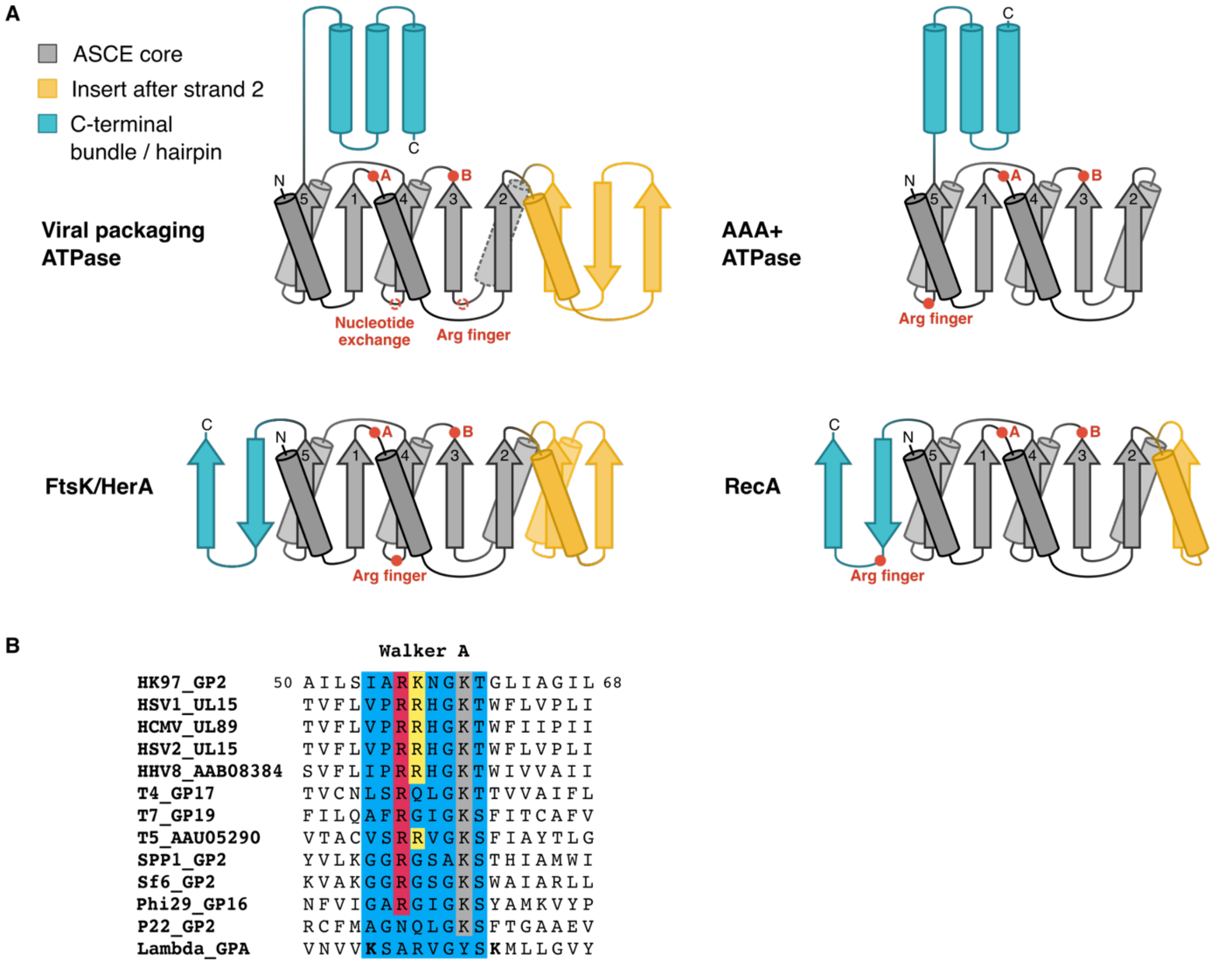


**Figure S3. Comparison of ASCE ATPase topologies and Walker A sequences**. (A) Topology diagrams of select ASCE (additional strand conserved E) ATPase subfamilies. AAA+, ATPases associated with diverse cellular activities. The consensus fold for viral packaging ATPases across all structurally characterized *cos*, *pac* and phi29 systems are shown. The gray helix in the ASCE core with a dotted outline is a random coil in the HK97 TerL. The phi29 ATPase has an additional β-strand between ASCE strand 5 and preceding helix (not shown). The location of confirmed *trans*-acting arginine fingers are indicated for each ATPase subfamily. (**B**) Multiple sequence alignment of viral packaging ATPase Walker A sequences. The virus and gene product names are indicated, separated by an underscore. In grey is the classical Walker A lysine; red, critical arginine residue identified in *pac* phage T4, *cos* phage lambda and phi29; yellow, critical lysine residue in HK97 and corresponding positive residues in phage T5 and herpes viruses. Lambda gpA has an unusual Walker A sequence with two lysine residues upstream and downstream of the classical position (in bold), both required for ATPase and DNA packaging activities (1, 2).

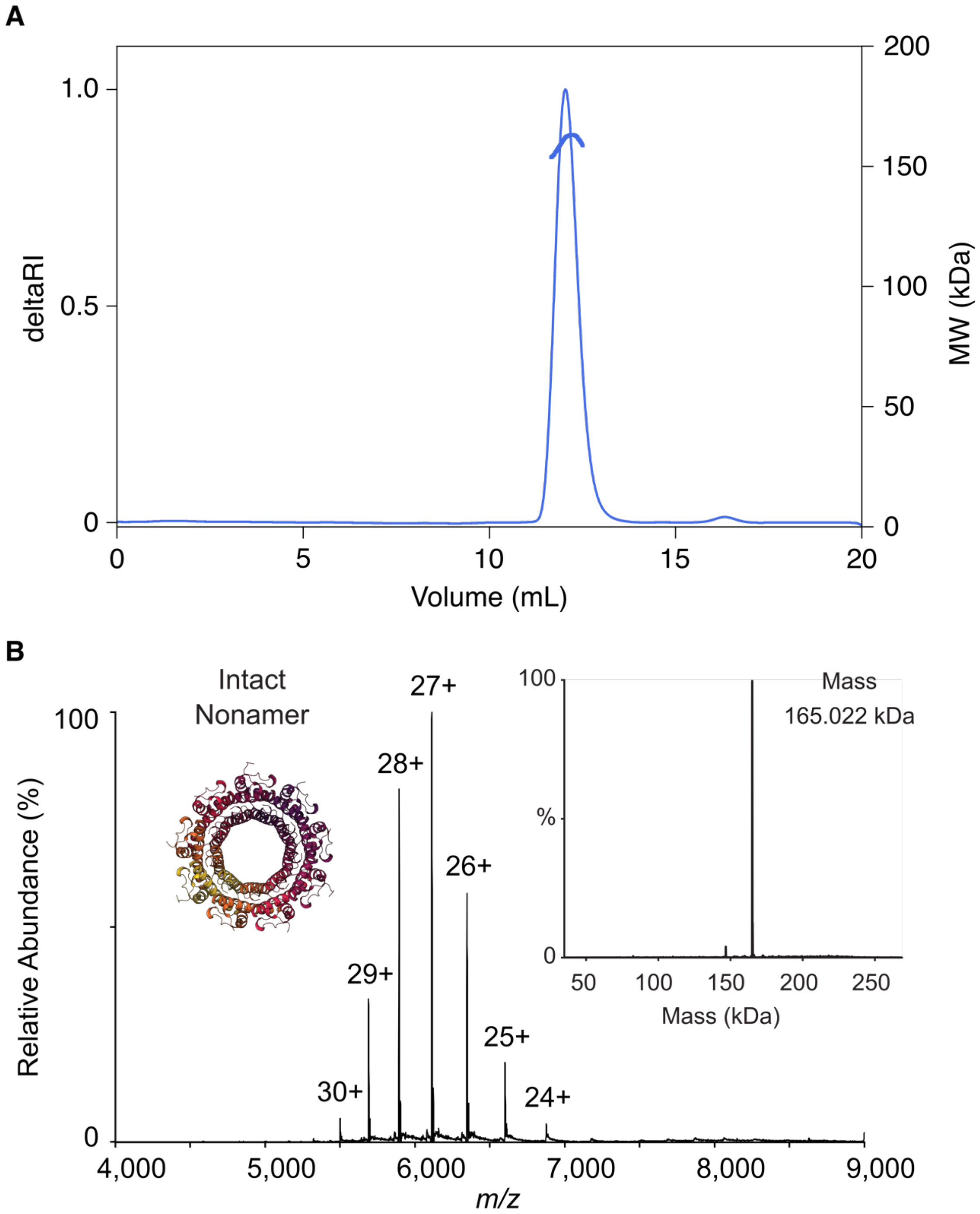


**Figure S4. Molecular weight determination of TerS.** (**A**) Size-exclusion chromatography and multi-angle laser-light scattering analysis (100 μL 2.7 mg/mL) in 20 mM HEPES, 300 mM KCl, 1 mM DTT, pH 7.0. Protein eluting from a Superdex 200 10/300 GL column (GE Healthcare) was analyzed using an on-line Dawn HELEOS-II light scattering detector and Optilab rEX refractometer (Wyatt). Data were processed based on a refractive index increment of 0.183 mL/g in ASTRA 5.3.4 (Wyatt). (**B**) High resolution Orbitrap native MS measurement showed a single charge state distribution corresponding to a species with a mass of 165.020±0.005 kDa (based on four most intense charge states). The fully deconvoluted spectrum indicated a single species with mass of 165.022 kDa (inset), unambiguously indicating a nonameric assembly based on the known monomer mass of 18.3 kDa.

**Table S2. TerS crystallographic data collection and refinement statistics.**

| | 500 mM KI soak | 250 mM KI soak |
|---|---|---|
| **Data collection** | | |
| Wavelength (Å) | 1.6984 | 0.9763 |
| Space group | *H*3 | *H*3 |
| Cell dimensions | | |
| *a*, *c* (Å) | 123.3, 79.4 | 123.3, 79.0 |
| Resolution (Å) | 44.31–1.79 (1.83–1.79) | 44.23–1.40 (1.43–1.40) |
| $R_{merge}$ | 0.080 (0.539) | 0.049 (0.707) |
| $I / \sigma(I)$ | 15.2 (1.4) | 10.3 (1.1) |
| $CC_{1/2}$ (%) | 99.6 (51.5) | 99.7 (52.5) |
| Completeness (%) | 95.3 (47.3) | 98.8 (88.6) |
| Redundancy | 4.4 (1.2) | 2.7 (2.0) |
| Wilson B ($Å^2$) | 19.0 | 18.6 |
| No. unique reflections | 40412 (1184) | 86871 (3870) |
| **Refinement** | | |
| Resolution (Å) | | 44.23–1.40 |
| No. reflections | | |
| Working | | 82402 |
| Free | | 4469 |
| $R_{work}/R_{free}$ | | 0.159/0.170 |
| No. atoms | | |
| Protein | | 2474 |
| Water | | 338 |
| Ligand | | 9 |
| B factors ($Å^2$) | | |
| Protein | | 21.9 |
| Water | | 33.0 |
| Ligand | | 30.9 |
| R.m.s. deviations | | |
| Bond lengths (Å) | | 0.011 |
| Bond angles (º) | | 1.4 |
| Ramachandran | | |
| Favored (%) | | 100.0 |
| Outlier (%) | | 0.0 |

**Table S3. Intramolecular contacts in the N-terminal helix-turn-helix-like region of HK97 TerS.**

| Residue 1 | Residue 2 | Nature of contact |
|---|---|---|
| T25 | H43 | hydrogen bond between sidechains |
| I26 | W42 | hydrogen bond between backbone oxygen and sidechain |
| P28 | W42 | CH/π stacking interaction |
| P29 | A32 | hydrogen bond between backbone atoms |
| S30 | G33 | hydrogen bond between backbone atoms |

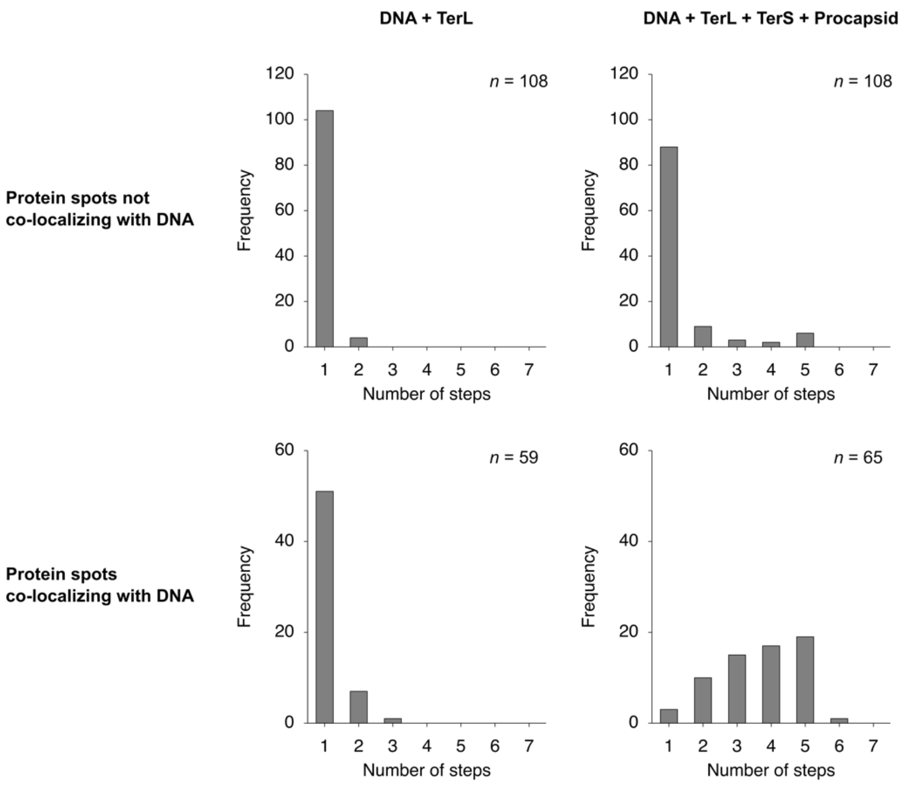


**Figure S5. GFP-TerL photobleaching steps observed by total internal reflection fluorescence microscopy.** Left column, GFP-TerL in the presence of only DNA; right column, GFP-TerL in the presence of DNA, TerS and procapsids. Top row, events not coincident with BOBO3 DNA staining. Bottom row, events coincident with DNA staining.

**Table S4. Event counts in the step-wise photobleaching assay.**

| | **DNA + TerL** | **DNA + TerL + TerS + Procapsid** |
|---|---|---|
| **Number of DNA spots** | **699** | **367** |
| with no co-localizing protein signal | 634 | 259 |
| with co-localizing protein signal, discarded | 6 | 43 |
| with co-localizing protein signal, analyzed | 59 | 65 |

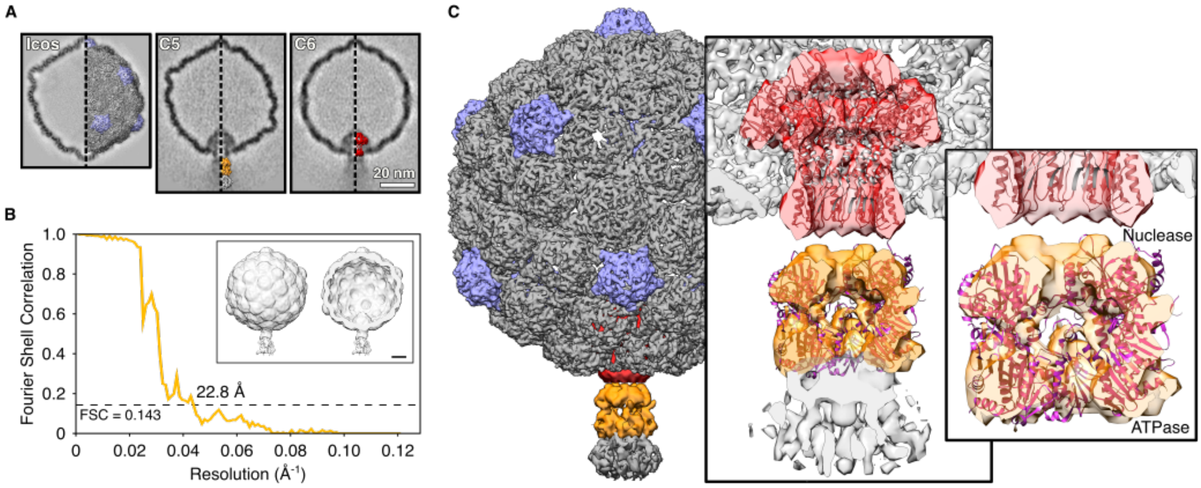


**Figure S6. Low-resolution cryo-EM-based model of the HK97 DNA packaging assembly**. (**A**) Cross-section of three reconstructions with icosahedral, C5 and C6 symmetry imposed. The poorly resolved densities corresponding to terminases and DNA indicated a dynamic system. 3D densities depicted on the right side of each cross-section were used to generate a composite map. (**B**) Fourier shell correlation analysis of the final half-maps during refinement with no symmetry imposed. Inset shows densities from this C1 refinement (whole and cut-through). Scale bar = 100 Å. (**C**) Composite map showing portal (red) and TerL (orange). Inset, rigid body fitting of a HK97 portal homolog (PDB 3KDR), and model of the HK97 TerL pentamer based on *pac* phage P74-26 pentameric ATPase model. It is unclear if the distal density corresponds to DNA or TerS.

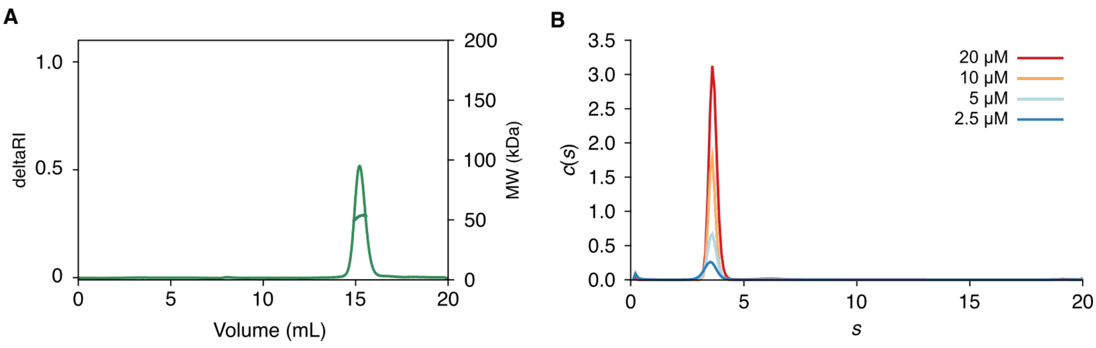


**Figure S7. Molecular weight determination of TerL.** (**A**) Size-exclusion chromatography and multi-angle laser-light scattering analysis (100 μL 1.2 mg/mL) in 20 mM HEPES, 300 mM KCl, 1 mM DTT, pH 7.0 using a Superdex 200 10/300 GL column (GE Healthcare). (**B**) Sedimentation coefficient distribution by sedimentation velocity analytical ultracentrifugation. Epon double-sector centerpieces were filled with protein (2.5-20 uM in 20 mM HEPES, 300 mM KCl, 1 mM DTT, pH 7.0) and buffer, respectively, and centrifuged in an An-60 Ti rotor at 50000 rpm, 20 °C, for 7.5 h using a Beckman Optima XL/I ultracentrifuge. Absorbance scans were acquired at 280 nm at 7.5 min intervals. A protein partial specific volume of 0.73747 mL/g, buffer density and viscosity of 1.01390 g/mL and 0.01002 P was estimated using Sednterp (3). Data were analyzed under the continuous c(*s*) distribution model using Sedfit (4). Frictional ratio estimates of 1.32–1.37 and standard sedimentation coefficients of 3.66–3.81 S were obtained, contributing to over 90% of the signal.

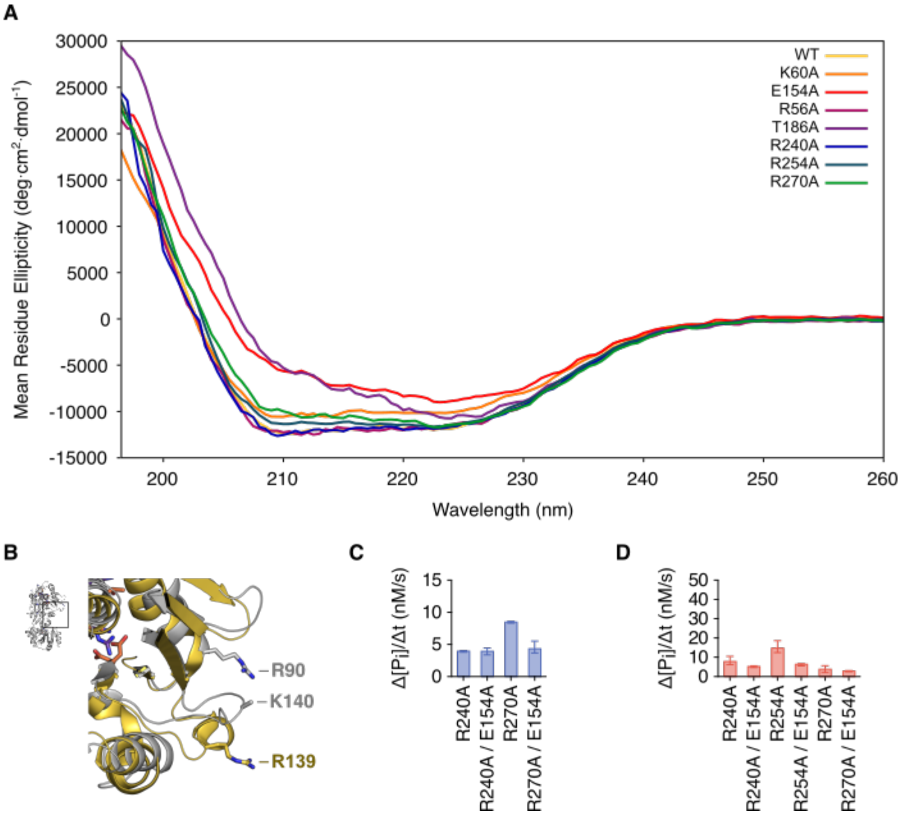


**Figure S8. Alanine mutagenesis analysis of TerL.** (**A**) Far-UV circular dichroism spectra of wild-type and mutant HK97 proteins. Spectra were recorded for protein (2–5 μM) in 20 mM Tris sulfate, 50 mM $Na_2SO_4$, 10 mM $MgSO_4$, 0.5 mM tris(2-carboxyethyl)phosphine hydrochloride, pH 7.5, at 20 °C. (**B**) Superposition of the HK97 TerL structure with the arginine finger of *pac* phage P74-26 TerL (gold, PDB 4ZNK) (**C**, **D**) ATPase rescue experiments conducted with TerL Walker B E154A mutant in equimolar amounts at 1 mM ATP concentration in the absence and presence of DNA, TerS and procapsids, respectively.

**Table S5. DNA oligonucleotides for cloning and mutagenesis.** Primers for plasmid linearization for restriction-free cloning are listed. Numbering on *cos* DNA primers indicates the genome position of the first complementary nucleotide. All sequences are provided in 5' to 3' direction.

| | |
|---|---|
| **Restriction-free cloning of His-SUMO-fusion** | |
| SUMO_GP1_for | gaacagattggtggtgcagataaacgaatccgttccg |
| SUMO_GP1_ rev | tacctaagcttgtctttatccgtgcttgggaaaggc |
| SUMO_GP2_ for | gaacagattggtggtatgacgcgaggtgagcg |
| SUMO_GP2_rev | tacctaagcttgtcttcacatgctcagcggtcg |
| Plasmid_for | agacaagcttaggtatttattcgg |
| Plasmid_rev | accaccaatctgttctctgtga |
| **Restriction-free cloning of His-fusion** | |
| His_GP2_for | atcaccaccaccacatgacgcgaggtgagcg |
| His_GP2_rev | tgaggagaaggcgcgtcacatgctcagcggtcg |
| Plasmid_for | cgcgccttctcctcactgttccaggggccccat |
| Plasmid_rev | tgtggtggtggtgatgatggctgctgccc |
| **Site-directed mutagenesis** | |
| GP2_R3A_for | cggcaggtgagcgtgtaatagcgttcatt |
| GP2_R3A_rev | ctcacctgccgtcatgtggtggtgg |
| GP2_R6A_for | gtgaggccgtaatagcgttcattgagcg |
| GP2_R6A_rev | tattacggcctcacctcgcgtcatgtgg |
| GP2_R13A_for | attgaggccttttgcatcgtgccagaag |
| GP2_R13A_rev | caaaaggcctcaatgaacgctattacacgc |
| GP2_R28A_for | ctatggcgttggacccctttcagaaagattt |
| GP2_R28A_rev | tccaacgccataggttggccgataagc |
| GP2_R56A_for | cgccgcgaaaaatggtaagactggcc |
| GP2_R56A_rev | tttttcgcggcgatggagaggatcgc |
| GP2_R90A_for | gcgcggaacaggcggccatcg |
| GP2_R90A_rev | tgttccgcgctgagtgcaccgct |
| GP2_R159A_for | ggttgcgggcccgcaggatgattttatc |
| GP2_R159A_rev | gcccgcaacctgccctgtttcatcg |
| GP2_R226A_for | agtaaagccgagtcctggctggctgc |
| GP2_R226A_rev | gactcggctttactgatatcagcgtcttttgg |
| GP2_R240A_for | cattcgcgtcagaaaaagacatggcgc |
| GP2_R240A_rev | ctgacgcgaatgttcccagtgccg |
| GP2_R247A_for | gcggcccaggctgagaaagctgg |
| GP2_R247A_rev | ctgggccgccatgtctttttctgacc |
| GP2_R254A_for | ctggcgcaatgccaagcttcgaaaacac |
| GP2_R254A_rev | cattgcgccagctttctcagcctggc |
| GP2_R263A_for | cttcgcgaacctcaacctcaatcagcg |
| GP2_R263A_rev | aggttcgcgaaggtgttttcgaagcttg |
| GP2_R270A_for | tcaggccgtgtctaccgtatcgccg |
| GP2_R270A_rev | acacggcctgattgaggttgaggtttcg |
| GP2_K140A_for | aaggtgcgacgacgcacggcc |
| GP2_K140A_rev | cgtcgcaccttctgcggataaagcc |
| GP2_K204A_for | gtcgcgtcgaaagatccgcacatcg |

| | |
|---|---|
| GP2_K204A_rev | ttcgacgcgaccgcatcatcaatccag |
| GP2_K206A_for | aatcggccgatccgcacatcgtgtg |
| GP2_K206A_rev | gatcggccgatttgaccgcatcatcaa |
| GP2_T186A_for | cagtgcgcaggcagcaaacgatgctga |
| GP2_T186A_rev | cctgcgcactgataacgattagcagcgg |
| GP2_K60A_for | ggtgcgactggcctgattgccggaat |
| GP2_K60A_rev | ccagtcgcaccatttttcgggcgatg |
| GP2_E154A_for | tcgatgcgacagggcaggttagggg |
| GP2_E154A_rev | ctgtcgcatcgagaatggccagaatgg |
| **GFP and linker insertion** | |
| GFP_for | atcaccaccaccacatggtgagcaagggc |
| GFP_rev | ccagatccgccgctgccacccttgtacagctcgtccat |
| Plasmid_for | cagcggcggatctggcggttccggtatgacgcgaggtgag |
| Plasmid_rev | tgtggtggtggtgatgatggctgctgccc |
| **Overlap-extension and restriction cloning of HK97 *cos* region** | |
| cos-312BamHIfor | gtggatccgacggtaagtttgttcagc |
| cos1strev | caaactttggcggcggtcatttg |
| cos2ndfor | gaccgccgccaaagtttgaatttaac |
| cos+472EcoRIrev | gtgaattcccgtgcatcacgatcg |
| **Generation of 5′ biotinylated DNA** | |
| cos-80for | atcaaatgagaatgaatcgcatc |
| cos+150rev | aaacctgcatgggacg-BIO |

**SUPPLMENTARY NOTE**

**Analysis of photobleaching events**

DNA particles were identified by extracting connected components from an average of the first 500 frames in the red channel. The intensity-weighted centroids of these components were taken to be DNA coordinates. Coordinates fewer than 10 px apart were rejected. For each coordinate in the red channel, given the translation vector from calibrating with TetraSpeck microspheres (ThermoFisher) emitting in both channels, the corresponding pixel in the green channel was found. A circle 3.5 px in radius, large enough only to enclose an event, was drawn around the center of this pixel. Mean intensity of the circle as a function of time was extracted and analyzed for photobleaching steps using the PIF algorithm (5) with Chung-Kennedy filtering (6). Over-fitted steps were identified by generating a counter-fit with steps between the original steps and comparing chi-squared statistics, as introduced by Kerssemakers et al. (7), and subsequently rejected.

For more robust analysis, a moving two-sample t-test (8) and a chi-squared minimization-based step-finding algorithm (7) were also applied. However, this resulted in a systematic under-fitting or over-fitting of steps, owing to the low signal-to-noise ratio of the experiment.

To build a distribution of the number of photobleaching steps per protein event co-localizing with DNA, an event was accepted only if steps could be fitted and if it had an intensity-weighted centroid within experimental error of the centroid of the DNA particle. A centroid-based measure was used because the density of events in the green channel was high and in turn background pixels surrounding each event could not be reliably defined for the fitting of Gaussian distributions. Adapting from Gelles et al. (9), experimental error was estimated to be the precision (two standard deviations) with which the x- and y-position of one GFP-LT monomer could be assigned relative to another, in time. We calculated one standard deviation to be 40.4 nm in both directions.

We note that the number of photobleaching events could be underestimated due to the presence of a dark GFP population as a result of misfolding or photobleaching prior to the experiment; and due to mis-fitting of two photobleaching events as one because the two events occurred within the same video frame, as described in detail by Ulbrich and Isacoff (10). The observed photobleaching rate of the GFP fusion protein was 0.10 $s^{-1}$, typical of GFP proteins (11). We also acknowledge the possibility of overestimating the number of photobleaching steps when the event monitored was coincident with other passively absorbed GFP molecules.

If one assumed all molecules observed in the DNA-bound motor experiment were pentamers, a binomial distribution with 72% probability of GFP being fluorescent could be fitted. However, it was likely that lower-order species as observed in the absence of prohead and small terminase also contributed to the distribution. Given the complexity of the experiment, it was reasonable to conclude only that the upper limit for the oligomeric state of TerL in a motor assembly was 5.
